# Myelin ensheathment and drug response of oligodendrocytes are modulated by stiffness of artificial axons

**DOI:** 10.1101/2023.08.11.552940

**Authors:** Mingyu Yang, Calliope J.L. Martin, Kavin Kowsari, Anna Jagielska, Krystyn J. Van Vliet

## Abstract

Myelination is a key biological process wherein glial cells such as oligodendrocytes wrap myelin around neuronal axons, forming an insulative sheath that accelerates signal propagation down the axon. A major obstacle to understanding myelination is the challenge of visualizing and reproducibly quantifying this inherently three-dimensional process *in vitro*. To this end, we previously developed Artificial Axons (AAs), a biocompatible platform consisting of 3D-printed hydrogel-based axon mimics designed to more closely recapitulate the micrometer-scale diameter and sub-kilopascal mechanical stiffness of biological axons. First, we present our platform for fabricating AAs with tunable axon diameter, stiffness, and inter-axonal spacing. Second, we demonstrate that increasing the Young’s modulus *E* or stiffness of polymer comprising the AAs increases the extent of myelin ensheathment by rat oligodendrocytes. Third, we demonstrate that the responses of oligodendrocytes to pro-myelinating compounds are also dependent on axon stiffness, which can affect compounds efficacy and the relative ranking. These results reinforce the importance of studying myelination in mechanically representative environments, and highlight the importance of considering biophysical cues when conducting drug screening studies.

## Introduction

Myelin is a lipid-rich membrane that ensheaths neuronal axons, thereby accelerating the efficiency of electrical signal conduction down the length of the axon^1,2^. Myelin is critical for the homeostatic function of mammalian nervous systems. Oligodendrocytes produce myelin in the central nervous system (CNS) and are fundamental to CNS development and myelin regeneration following injury^3,4^. During development, oligodendrocytes derive from oligodendrocyte progenitor cells (OPCs), which are maintained throughout adulthood and account for approximately 2-8% of the cells in the adult CNS^5^. Oligodendrocytes and OPCs inhabit a dynamic microenvironment within the CNS, which provides both biochemical and biophysical cues that regulate oligodendrocyte function. A growing body of evidence demonstrates that oligodendrocytes and OPCs are sensitive to external mechanical cues^6^ including ECM stiffness (and more broadly the stiffness of the material to which OPCs adhere)^7–11^, mechanical strain^12^, macromolecular crowding, and physical confinement^13,14^.

The sensitivity of OPCs to biophysical cues may play important roles in myelin pathology and neurodegenerative diseases. For example, neuroimaging data from human patients show correlations between neurological disease states, such as progressive multiple sclerosis (MS), and changes in the structural integrity and mechanical properties of brain tissue^15,16^. These findings are supported by murine models of demyelination, in which the Young’s modulus *E* of brain parenchyma decreased in response to acute demyelination from 240 Pa to 120 Pa, signifying a stiffness reduction correlated with disease progression^17,18^. In addition to tissue stiffness, other biophysical cues such as axon diameter and spacing may play important roles in oligodendrocyte biology and pathology. Under healthy conditions, axon diameter in the CNS can vary in diameter from 0.1-5.0 µm^19,20^. Magnetic resonance imaging (MRI) data has shown axon swelling to be a major pathological feature in chronic MS^21^, and a similar case of increased axonal diameter was reported for amyotrophic lateral sclerosis (ALS)^22^. Furthermore, axonal injury and loss of axon density (number of axons per unit area or volume) is a hallmark of progressive MS^23^, especially for chronically demyelinated axons. An open question is whether these biophysical changes simply act as correlative biomarkers or contribute to disease progression. In other words, do changes in axon diameter and brain parenchyma stiffness occur as secondary byproducts of myelination pathology, or do they actually change the propensity of oligodendrocytes to myelinate?

One approach for exploring these questions is to recapitulate and control these cues in an *in vitro* model. Prior examples of cell-free physical models of axons include electrospun fibers and glassy polymers. Although these models can recapitulate the geometry of biological axons, they use materials with Young’s modulus in the megapascal and gigapascal range, several orders of magnitude stiffer than biological axons^14,24–29^. With such materials, it is not possible to explore how changes in Young’s moduli within the sub-kilopascal physiological range can modulate myelination; nanometer-scale diameters and spacing of those stiff fibers can also challenge optical image-based validation of myelin wrapping around the fibers. To that end, we have previously developed Artificial Axons (AAs), which are 3D-printed hydrogel structures that mimic the sub-kilopascal stiffness and micrometer-scale diameter of biological axons^30–32^. Importantly, in this platform we can engineer properties of the axon arrays, such as stiffness, spacing, and diameter, enabling systematic studies of the influence of each cue on myelin wrapping. OPCs seeded on the AAs differentiate into mature oligodendrocytes and ensheath the axons with myelin, which we quantify in 3D using an in-house image analysis pipeline.

While the AAs reported previously were a demonstrated proof of concept, scalability was limited and three-dimensional (3D)-printing of AAs on individual coverslips required at least two weeks to create sufficient samples for testing (for instance, to fill a 96-well plate). Here we describe advancements in design and fabrication of the AA platform, in which we 3D-print AAs directly into a 96-well plate using a digital mask, producing AAs with independently tunable Young’s modulus, axon diameter, and inter-axonal spacing. With this new platform, we eliminated sample damage to increase yield and reduced the total fabrication time to two hours, enabling production of a 96-well plate in which each well could represent a distinct combination of axon stiffness, diameter of spacing. We leveraged this tunability to explore how each parameter affects myelin ensheathment by rat oligodendrocytes. Finally, we focused specifically on Young’s modulus and investigated how axon stiffness influenced oligodendrocyte responses to pro-myelinating compounds. We found that the relative efficacy of different pro-myelinating compounds, as quantified by a myelin wrapping index, differed between the AAs of higher (*E* = 13000 Pa) and lower stiffness (*E* = 800 Pa) axons, with different ‘top 3’ drug rankings identified in each condition. In summary, we demonstrate that the AA platform is amenable to quantifying correlative and discovering causal relationships between axon biophysical cues and myelin ensheathment by oligodendrocytes, and that the mechanical environment can influence oligodendrocytes’ response to soluble biochemical cues including pro-myelinating compounds.

## Materials and Methods

### Ethics Statement

This study was carried out in accordance with the guidelines of the National Institutes of Health for animal care and use (Guide for the Care and Use of Laboratory Animals) and the protocol was approved by the Institutional Animal Care and Use Committee at the Massachusetts Institute of Technology (MIT Committee on Animal Care).

### Fabrication of Artificial Axons (AAs)

Artificial axons were fabricated directly in 96-well plates, using a projection microstereolithography setup. To vary axon stiffness (material Young’s modulus) the resins were prepared with varying ratios of 1,6-hexanediol diacrylate (HDDA) (Sigma-Aldrich) and 4-arm PEG acrylate (starPEG) (JenKem) monomers, at mass ratios of 3:1, 2:1, and 1:1. The resins were pipetted into the wells of 96-well plates and exposed to UV light using our custom made projection microstereolithography setup^31^. Projecting the UV light onto the resin through the digital mask caused the liquid monomer resin to polymerize into solid vertical pillars that match the geometric pattern of the mask.

The shape of the mask, the composition of the resin, the UV exposure duration, and the UV exposure intensity could all be modified to independently tune the Young’s modulus, diameter, and spacing of the axons. For convenience, for the experiments on drug-induced myelination on different stiffness axons, we used a physical mask with the area corresponding to the area of the entire plate bottom, to generate axons in all plate wells simultaneously. Prior to introduction of rat oligodendrocyte progenitor cells, the AAs were functionalized with poly-D-ornithine (Sigma-Aldrich) (50 µg/mL) followed by incubation with laminin (Gibco) (20 µg/mL) to facilitate cell adhesion. The completed AA plates were stored in PBS at 4°C and warmed to 37°C the day of OPC seeding.

### Mechanical characterization of AAs

The Young’s modulus *E* of the cured AA material was determined using atomic force microscope (AFM)-enabled nanoindentation measurements (MFP-3D Bio, Asylum Research). Cylindrical structures of each material (10 µm thickness and width) were fabricated by projection microstereolithography using the same printing conditions as the AAs, and equilibrated overnight in PBS. AFM measurements were performed using a cantilever with nominal spring constant *k* = 0.1 N/m, terminating in a poly(methyl methacrylate) spherical probe with approximate diameter 1.5 µm (NanoAndMore). The actual spring constant was calibrated via the thermal noise method^33^. Between 30 and 40 force-depth responses were collected from each sample of the material. The cantilever base velocity was 1 µm/s, and probe retraction was triggered after reaching a maximum force of 30-100 nN, with lower forces for the more compliant samples. Young’s moduli *E* were calculated by fitting the spherical Hertzian elastic contact model^34^ for data acquired up to an indentation depth of 200 nm.

### 3D myelin wrapping assay

Rat oligodendrocyte progenitor cells (rOPCs) were isolated from neonatal rat brains (postnatal day 1) using magnetic sorting with beads coated with A2B5 antibodies (Miltenyi). The isolated cells were expanded in tissue culture flasks for 2-3 days in proliferation medium containing DMEM/F-12 media (Gibco), penicillin-streptomycin (Gibco), B-27 (Gibco), 10 ng/mL each of platelet-derived growth factor (PDGF) (Gibco), and fibroblast growth factor (FGF) (Gibco). Expanded rOPCs were seeded in 96-well AA plates at a density of 20,000 cells per well in differentiation medium, consisting of DMEM/F-12 media, B-27, and 2 ng/mL each of PDGF and FGF. Only the inner 60 wells were used to avoid the outermost perimeter of wells which had accelerated liquid evaporation. 24 hours after seeding, 1/3 of the media was changed with fresh differentiation medium supplemented with a pro-myelinating drug. As the control condition, we used medium containing 0.1% DMSO, which was the solvent vehicle used for other compounds. Media exchange of 33% volume occurred every other day, and cells grew on the AAs for either seven or 14 days before being fixed and stained. Every condition was repeated in at least triplicate, and two independent biological replicates (separate rounds of cell culture with two different rOPC batches) were conducted.

### Immunostaining

Cells were fixed with 4% paraformaldehyde (Electron Microscopy Sciences), washed three times with PBS, and permeabilized with 0.1% v/v Triton X-100 and 5% v/v goat serum in PBS for 10 minutes at room temperature. Then, cells were washed three times in PBS and blocked for 1 hour in 5% v/v goat serum in PBS for 1 hour at room temperature. Cells were incubated in primary antibody (rat anti-MBP, BioRad, 1:200 dilution) for 24 hours at 4°C. Next, cells were washed three times in PBS and incubated with secondary antibody (Alexa-Fluor-647 goat anti-rat, Thermo Fisher, 1:200 dilution) for 1 hour at room temperature. Cells were then washed three times in PBS and incubated with DAPI (Thermo Fisher, 1:1000 dilution) for 5 minutes. Finally, cells were washed once more and stored in PBS at 4°C.

### Fluorescence imaging

Stained samples were imaged under three fluorescent channels (Alex-Fluor 647 for MBP+ myelin, rhodamine for AAs, DAPI for nuclei) using a confocal microscope (Olympus, FluoView 3000) at 20x air lens. For each well of the 96-well plate, eight fields of view were imaged; at each one, a confocal stack image was taken consisting of 8 *z-* slices separated by a step size of 2 µm. Collectively, ∼10,000 AAs were imaged and analyzed per well.

### Quantification of 3D myelin wrapping on AAs

The fluorescent optical *z*-slice images were processed through an in-house image analysis pipeline. In brief, the AA, MBP+, and DAPI channels were imported to ImageJ and thresholded to obtain binary masks, followed by 3D volume reconstructions of the *z*-stacks. A 1-pixel-thick outline was traced around each AA; the outline was compared to the myelin mask to quantify the fraction of each axon circumference ensheathed by myelin. Aggregating across all *z*-slices, AA pillars were classified as ‘fully wrapped’ if they exhibited a contiguous >6 µm ensheathed segment length in which the AA pillar was >80% wrapped in oligodendrocyte-synthesized myelin membrane positive for myelin basic protein (MBP) across all *z*-slices of that pillar height. For each field of view, a myelin wrapping index (WI) was calculated, as the number of fully wrapped artificial axons normalized by the number of cell nuclei.

### Statistical analysis

All imaged fields of view from a given experimental condition were pooled and averaged to determine a mean wrapping index per condition. For pairwise comparisons between conditions, Mann Whitney Wilcoxon tests were performed using the SciPy package in Python. For three-way comparisons between three tested conditions, Kruskal-Wallis tests were performed using the SciPy package. For drug response experiments, pairwise comparisons between conditions were done using one way ANOVA with Bonferroni correction within Origin Pro software.

## Results and Discussion

### Additive manufacturing of Artificial Axons (AAs)

We fabricated Artificial Axons using projection microstereolithography, a 3D-printing approach that uses UV light to polymerize micrometer-scale solid structures. We previously developed a custom poly(HDDA-*co*-starPEG) resin which is liquid when unpolymerized, but solidifies into a columnar geometry when exposed to UV light^30–32^. Our previously reported approaches utilized individual 5 mm-diameter coverslips to fabricate AA arrays, which had limited scalability and required at least two weeks to create sufficient samples for testing in a full 96-well plate. Furthermore, the delicate sample handling requirements led to low yield of useful coverslips, with disproportionate damage of the samples with AAs of lower diameter (<10 µm) and lower stiffness (<1 kPa).

**Figure 1A** demonstrates a new fabrication approach in which we polymerize AAs directly into a multiwell plate using digital photomask, which consists of an array of white circles on a black background. The digital mask modifies the UV light incident upon the resin, thus giving rise to vertical axons that match the pattern of the mask. Modification of the parameters of the digital mask via custom LabView code provides the flexibility to tune the AA geometry on a per-well basis by applying distinct masks and UV light exposure time and intensity. This approach reduced the total fabrication time of a 96-well plate of AA arrays from two weeks to two hours. **Figure 1A** shows the potential to independently vary axon diameter and inter-axonal spacing using the digital photomask approach, simply by varying the size and spacing of white dots in the digital mask.

**Fig. 1:**
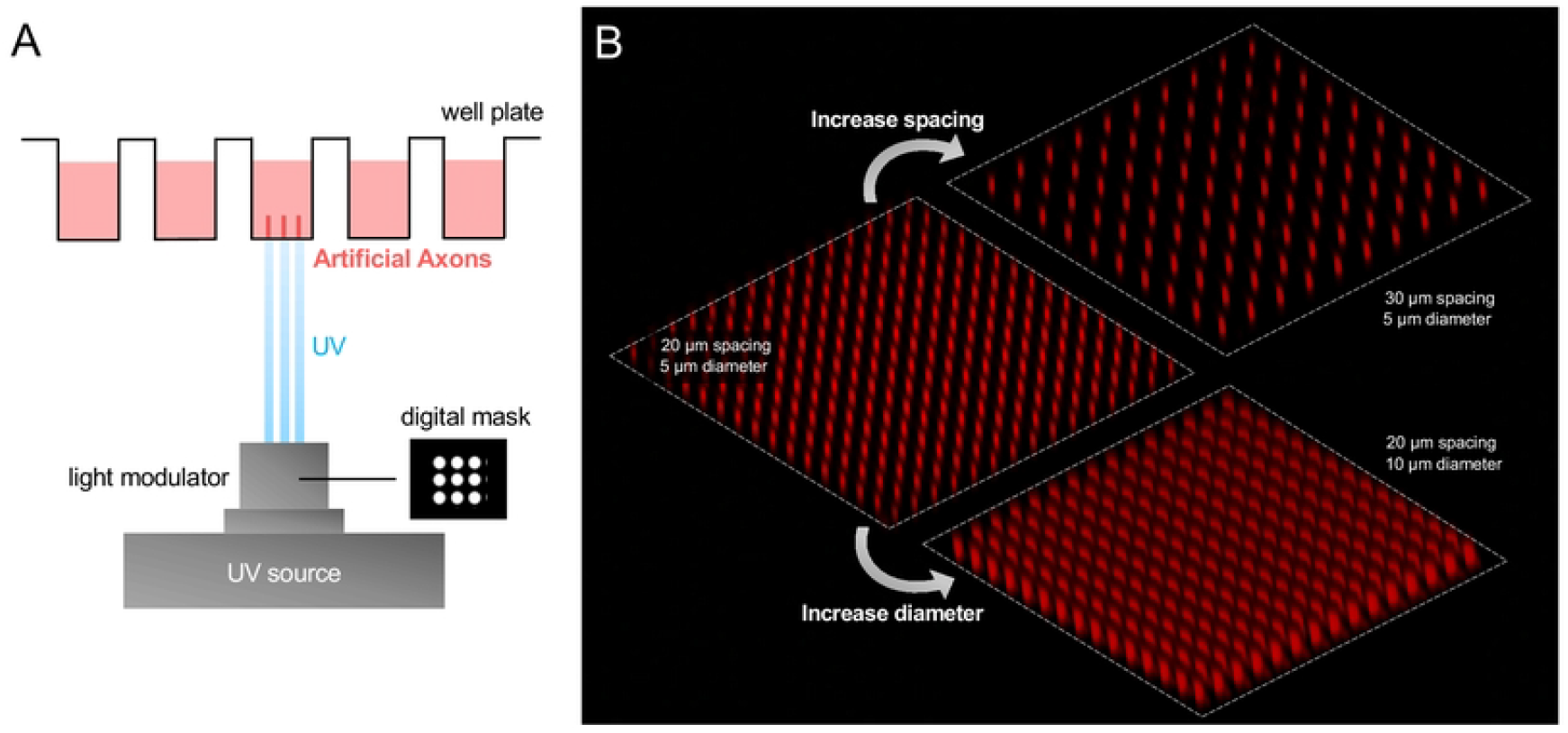
(A) Schematic of Artificial Axon fabrication using a digital photomask. (B) Confocal micrographs of Artificial Axons with tunable spacing and diameter.

### Fine-tuned control of AA stiffness and diameter

The projection microstereolithography approach for manufacturing AAs provides multiple knobs for tuning AA properties, including the monomer chemistry and UV exposure conditions. **Figure 2A** shows the two primary components of our AA material: 4-arm PEG acrylate (starPEG) and 1,6-hexanediol diacrylate (HDDA). Copolymerization of starPEG and HDDA forms a network polymer illustrated in **Fig. 2B**. We combined HDDA and starPEG at three different mass ratios of 3:1, 2:1, and 1:1 and determined their Young’s moduli *E* using atomic force microscope (AFM)-enabled indentation (see Materials and Methods). In order of descending HDDA:starPEG ratio, the *E* of the three materials were found to be 13000 Pa ± 64 Pa, 780 Pa ± 11 Pa, 98 Pa ± 5 Pa, respectively. This trend was consistent with expectations because HDDA functions as a crosslinker between the much larger starPEG molecules in the polymer network. Therefore, increasing the HDDA:starPEG ratio represents an increase in crosslinking density and therefore anticipates a stiffer polymerized AA material. Importantly, Materials Y and Z are consistent the range of stiffnesses reported for neural tissue and axons.

**Fig. 2:**
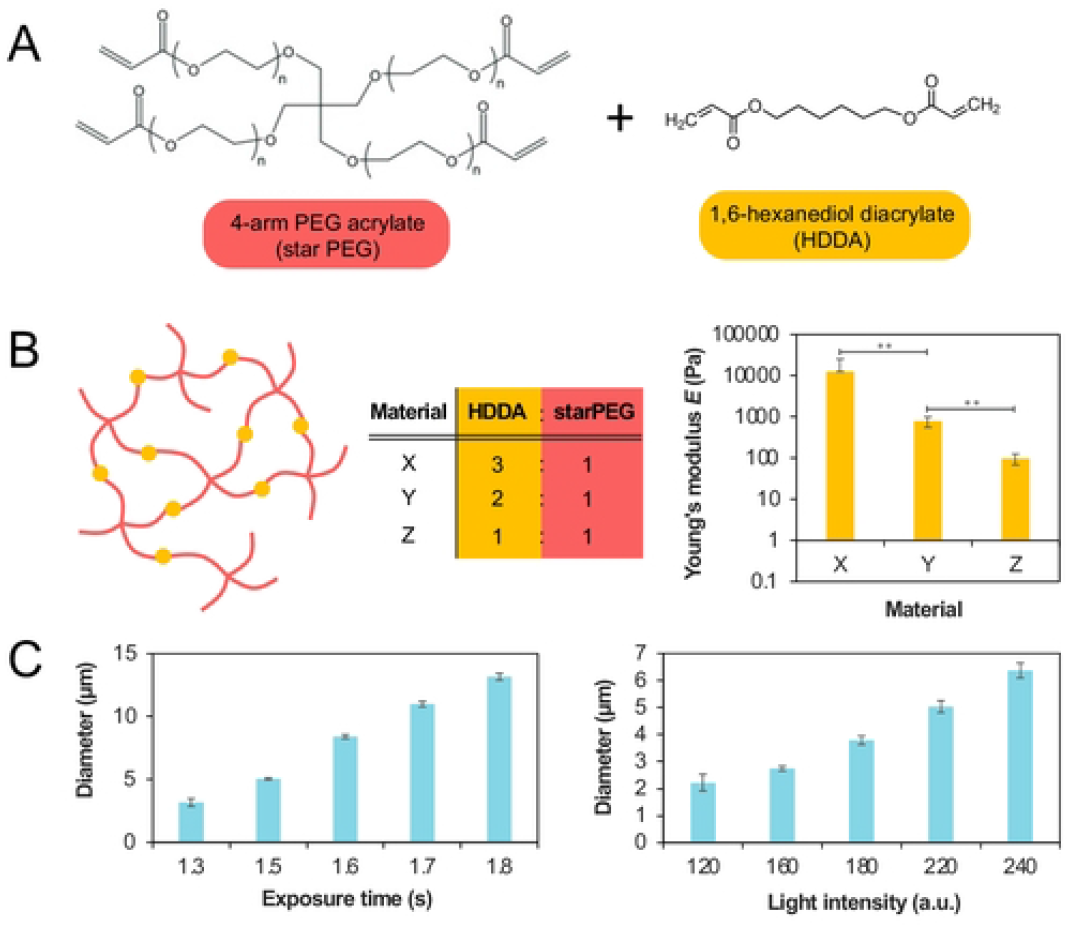
(A) Chemical structure of 4-arm PEG acrylate and 1,6-hexanediol diacrylate, the two main components of Artificial Axons. (B) Schematic of network structure formed by UV-mediated crosslinking of starPEG and HDDA. Materials X, Y, and Z are defined as having a 3:1, 2:1, and 1:1 HDDA:starPEG ratio respectively. Young’s elastic modulus (*E*) measured for three Artificial Axon samples with different HDDA:starPEG weight ratios. Error bars represent standard error of the mean. (C) Tunability of AA diameter by varying the UV exposure time and light intensity. Light intensity is measured on an arbitrary scale from 0 to 255. Data shown for Material X, and error bars represent standard error of the mean.

To investigate the role of UV exposure time and intensity on AA diameter, we produced a digital photomask comprising an array of single-pixel white dots on a black background. We projected a UV beam of this mask to polymerize Material X (3:1 HDDA:starPEG ratio), across a range of exposure times (0.5s – 2s) and light intensities (100-250, measured on an arbitrary scale of 0-255). Finally, we used confocal microscopy and ImageJ^35^ to quantify the distribution of resultant AA diameters. As expected, increasing both the UV exposure time and light intensity produced a concomitant increase in axon diameter. This is consistent with the free-radical mediated polymerization mechanism through which the AAs form. UV exposure initiates polymerization by imparting reactive free radicals onto the monomer species. Increasing the light intensity increases the quantity of free radicals produced, thereby causing more monomers to react and producing larger-diameter axons from the same 1-pixel digital mask. A similar argument holds for increasing the exposure duration. Furthermore, we found that UV exposure duration and intensity both had to exceed a critical threshold for polymerization (i.e., in order for any AAs to form). In sum, adjusting the monomer chemistry and exposure conditions enables fine control over both axon stiffness and diameter, thus paving the way to investigate how these parameters may influence myelination by oligodendrocytes.

### Influence of AA stiffness, diameter, and density on myelin wrapping by rat oligodendrocytes

We fabricated AAs of three different magnitudes of stiffness, of diameter, and of spacing, and quantified the resulting myelin ensheathment by rat oligodendrocytes as a parameter we termed wrapping index, WI. OPCs were isolated from neonatal rat brains and seeded on AAs within 96-well plates for 14 days with differentiation induced by T3 at concentration 1 µM, after which we fixed and stained for myelin basic protein (MBP), a marker of differentiated oligodendrocytes and also a major component of myelin itself. **Figure 3A** shows a schematic of our in-house image analysis pipeline for quantifying 3D myelin ensheathment (see Materials and Methods). In brief, we used confocal microscopy to take multiple top-down *z*-slices of the myelin-ensheathed AAs, then used ImageJ to compare across *z*-slices to quantify myelin wrapping. Aggregating across all *z*-slices, we classified pillars were classified as fully wrapped if they had a contiguous >6 µm ensheathed myelin segment in which the AA was >80% wrapped around the AA pillar circumference across all *z*-slices. For each field of view, we calculated the wrapping index WI, defined as the number of fully wrapped axons normalized by the number of cell nuclei. This allowed us to compute a mean wrapping index for each AA stiffness, spacing, and density condition.

**Fig. 3:**
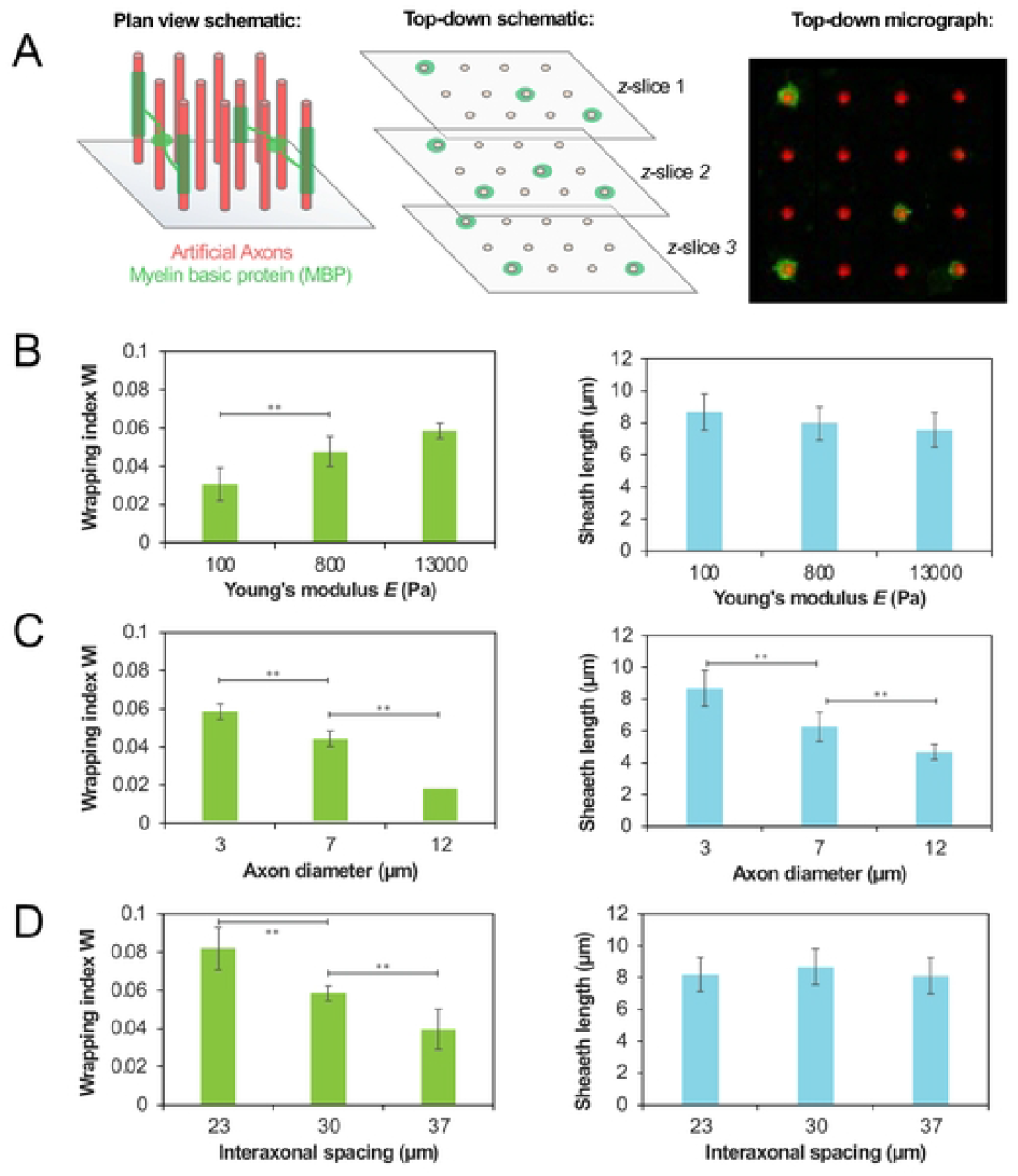
(A) Schematic of oligodendrocytes depositing myelin basic protein (MBP)-containing myelin on AAs. These are visualized through top-down confocal micrographs taken across multiple *z*-slices. Scalebar in micrograph is 30 µm. Variation of oligodendrocyte myelin wrapping with (B) Young’s modulus, (C) diameter, and (D) density. The left graph shows wrapping index WI, a measure of the *number* of AAs wrapped. The right graph shows the distribution of myelin sheath lengths on AAs. The data for panels C and D are for the *E* = 13000 Pa axons. Error bars represent standard error of the mean.

**Figures 3B-D** show the variation of myelin wrapping by oligodendrocytes as a function of AA stiffness, diameter, or spacing. Importantly, we measured two attributes of myelin wrapping. The first is the previously defined WI, which indicates the *number* of AAs wrapped. We also separately quantified the *length* of the imaged myelin sheaths, calculated by determining the number of adjacent *z*-stacks with >80% wrapping. The data shown in **Fig. 3** shows one biological repeat of the experiment; the data for a second replicate repeat are shown in **Supplementary Fig. 1**.

**Figure 3B** shows that increasing axon stiffness led to an increase in the WI, meaning that on average more AAs are being wrapped per OPC (left panel). The average length of myelin segment shows slightly decreasing trend with the increasing stiffness, although the differences between the individual tested conditions were not statistically significant (right panel). This raises the possibility that the decrease in stiffness local to OPCs or oligodendrocytes in demyelinating contexts^15,17^could possibly decrease the intrinsic propensity of oligodendrocytes to myelinate axons. Based on these data alone, this is still just a speculative hypothesis; furthermore, an important caveat is that in this experiment we vary stiffness on an *axon*, whereas previous data report on more global changes in stiffness of the brain *tissue* in demyelinating lesions, without capacity for delineating stiffness at the individual cell or axon level.

**Figure 3C** shows that increasing AA diameter led to a decrease in WI. Furthermore, the myelin that *was* wrapped on the higher-diameter AAs on average had shorter lengths. We note that the data reported in literature for biological axons or axon mimicking fibers with diameters below 2µm demonstrates that within that range the larger-caliber axons are preferentially myelinated^36^, whereas our data consider axon mimics with diameters larger than 3 µm, which can model swollen axons in the inflammatory demyelinating lesions (and are also relevant to the PNS axon diameters). We continue further refining of our platform to capture sub-2µm axon diameter variation, closer to the biological range of the CNS axon diameters. **Fig. 3D** shows that increasing the mean separation between axons (in other words, reducing the axon density), which can model decreased axon density in chronic lesions, also decreased the WI, although with minimal effect on sheath length. In summary, these data show that AA stiffness, diameter, and spacing can all influence myelin wrapping by oligodendrocytes principally by affecting the *number* of AAs wrapped (captured by the WI), although axon stiffness and diameter can also influence the *lengths* of the myelin sheaths deposited on the AAs by maturing oligodendrocytes.

### Influence of AA stiffness on response to pro-myelinating compounds

Based on the observed influence of axon stiffness on myelin wrapping (**Fig. 3B**), we hypothesized that the responses of oligodendrocytes to pro-myelinating compounds may also depend on axon stiffness. To test this hypothesis, we conducted myelin wrapping assay in the presence of several pro-myelinating compounds acting on various ligands and signaling pathways, on axons with two distinct stiffnesses, 13 kPa (material X) and 0.8 kPa (material Y), with the AAs of lower stiffness corresponding approximately to that of normal axons^37^. Rat oligodendrocytes where cultured for 7 days and dosed every other day with a compound at concentrations of 3 µM (ketoconazole, clemastine, benztropine, quetiapine, clobetasol, fasudil and miconazole) or 100 nM (tasin-1, tamoxifen, amorolfine, bazedoxifene, and T3). The chosen drug concentrations corresponded to the maximum efficacy (measured as WI) in our previously conducted experiments on material X. At day 8, we fixed the cells and immunostained for MBP. We performed two independent experiments, each in triplicate.

**Figure 4A** shows that for many but not all tested compounds, the WI differed in magnitude on axons with different stiffness. We observed higher WI on stiffer axons upon oligodendrocyte exposure to ketoconazole, T3, quetiapine, clemastine, and benztropine. By contrast, responses for bazedoxifene and amorolfine were weaker (i.e., lower WI) on stiffer axons compared to those corresponding to physiological stiffness of axons. We did not observe statistically significant differences as a function of AA stiffness in responses to tasin-1 and tamoxifen, or for clobetasol, fasudil and miconazole for which WI was low overall and did not exceed the levels for the DMSO negative control condition. Notably, oligodendrocyte responses to amorolfine included WI significantly exceeding the DMSO response for axons of physiological stiffness, but insignificant wrapping above the negative control on stiffer axons. Conversely, several compounds inducing significant wrapping activity on stiffer axons (clemastine, benztropine, miconazole) showed negligeable activity on more compliant axons. The data in **Fig. 4A** represent the average of two biological replicates; the individuated data from each replicate are shown in **Supplementary Fig. 2**.

**Fig. 4:**
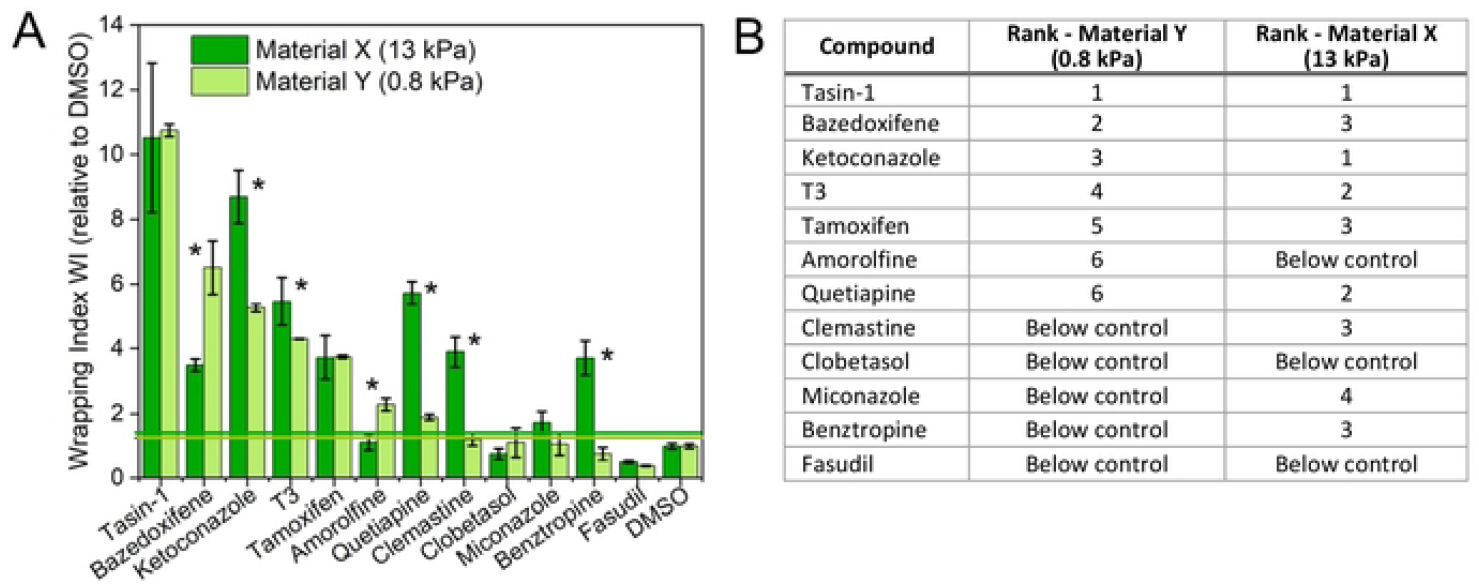
(A) Effect of stiffness on promyelinating activity of compounds. (A) Wrapping index WI for compounds scaled by control (WI for DMSO) on stiffer (Young’s modulus 13 kPa, dark green) and more compliant axons (Young’s modulus 0.8 kPa, light green). Error bars are standard error of the mean. (*) statistically significant difference between responses on materials X and Y, p < 0.05. Horizontal lines denote 2 standard deviations above the respective DMSO controls. (B) Rank order of WI for compounds can differ on more compliant (0.8 kPa) and stiffer (13 kPa) axons. Materials X and Y are HDDA-starPEG hydrogels differing in extent of crosslinking resulting in different material stiffness.

Additionally, relative ranking of WI among these compounds (**Fig. 4B**) differed when assessed on stiffer compared to more compliant axons (**Fig. 4B**). While the compound eliciting the highest WI under these dosages and conditions was the same for both AA stiffnesses (tasin-1), the compounds ranked second and lower varied and could only be distinguished statistically for the more compliant axons. This indicates that using assay with axons with significantly higher stiffness than the physiological values, could result in either over- or underestimating of compounds efficacy and some promising compounds could be missed. This supports the concept that evaluation of pro-myelinating potential should include assays with mechanical stiffness representative of the target environment. We note that the above results reflect specific combination of biophysical and biochemical environment (specific axon coating, axon stiffness range, concentrations of compounds, cell batch), and we do not claim generalizable results for all conditions or compounds. Rather, we consider this example a proof-of-concept of the importance of the mechanically correct environment for drug screening.

## Summary

We demonstrate that the fabrication and implementation of 3D artificial axons can enable direct measurement of how biophysical cues influence the 3D process of axon engagement and wrapping, or myelin ensheathment, by rat oligodendrocytes. By modulating the material composition and fabrication conditions, we developed AAs with tunable Young’s moduli or mechanical stiffness (from the Pa to the kPa range), diameters (3 µm-15 µm) and axon densities. Increasing the Young’s moduli or density of AAs corresponded in a concomitant increase in the mean *number* of AAs ensheathed by oligodendrocytes as quantified by a myelin WI that was normalized for the total cell number, with minimal impact on the resulting sheath segment lengths. Increasing the AA diameter corresponded to a decrease in the mean number of AAs ensheathed *and* a decrease in the mean segment lengths. Although future work is needed to even more closely represent physical cues of physiological axons (for instance, fabricating AAs with sub-3 µm diameter), these results demonstrate capacity to probe correlative and causal relationships between biophysical cues and myelin wrapping *in vitro*.

An important application of *in vitro* modeling is the potential to compare and identify drug compounds that could stimulate myelin repair in demyelinating diseases. We found that the relative efficacies of pro-myelinating compounds differed depending on AA stiffness, as quantified by the wrapping index, which may have implications for *in vitro* drug screening. For example, the compound candidates identified in the *in vitro* drug screens with oligodendrocytes cultured on high-stiffness substrata such as tissue-culture polystyrene (TCPS) or stiff nanofibers may not be representative of the 3D wrapping responses predicted in a more compliant biomechanical environment. As the **Figure 4** suggests, it is possible that drugs which *may* have been ranked highly in mechanically compliant environments could be missed when screening oligodendrocyte response in formats of superphysiological stiffness. Collectively, these results speak to the importance of studying myelination in mechanically representative environments.

## Acknowledgements

M.Y., A.J., and K.J.V.V. acknowledge funding from the U.S. Department of Defense Congressionally Directed Medical Research Program (MS190075/W81XWH2010365), the Deshpande Innovation Center at the Massachusetts Institute of Technology, and Sanofi. K.J.V.V. gratefully acknowledges the Michael (1949) and Sonja Koerner Professorship. M.Y. acknowledges the Angela Leong Fellowship Fund 2021-2022 and Hugh Hampton Young Memorial Fellowship 2022-2023 from the Massachusetts Institute of Technology.

## Authors Contributions

M.Y. and A.J. designed the research, conducted the experiments, analyzed and interpreted the data. K.K. led the design, construction, and optimization of the customized hardware and software components to develop the AA fabrication system. C.J.L.M. optimized the 3D-printing process and contributed to the data collection and analysis in Fig. 2. K.J.V.V. designed the research, helped interpret the data, and provided overall leadership of the project. M.Y., A.J., and K.J.V.V wrote the manuscript; all authors contributed to editing of the manuscript.

## Conflict of Interest

A. J. and K. K. are the founders and stock holders of Artificial Axon Labs Inc. K. J. V. V holds scientific advisor positions at Artificial Axon Labs. A. J. and K. J. V. V., are inventors on patents #US 20170328888A1 and #US 15/975,452.

